# The Amyloid Clearance Defect in ApoE4 Astrocytes is Corrected by Epigenetic Restoration of NHE6

**DOI:** 10.1101/243097

**Authors:** Hari Prasad, Rajini Rao

## Abstract

The accumulation of amyloid protein Aβ in senile plaques is a key driver and hallmark of Alzheimer disease (AD), a major cause of death and dementia in the elderly. The strongest genetic risk factor in sporadic AD is the ε4 allele of Apolipoprotein E (ApoE4), which potentiates pre-symptomatic endosomal dysfunction and defective clearance of Aβ, although how these two pathways are linked has been unclear. Here, we show that aberrant accumulation of endosomal protons in ApoE4 astrocytes traps the LRP1 receptor in non-productive intracellular compartments, leading to loss of surface expression and Aβ clearance. Hyperacidification of endosomal pH is caused by selective down regulation of the Na^+^/H^+^ exchanger NHE6, which functions as a critical proton leak pathway, in ApoE4 brain and astrocytes. *In vivo*, the NHE6^KO^ mouse model shows elevated Aβ in the brain. Epigenetic restoration of NHE6 expression with histone deacetylase inhibitors normalized ApoE4-specific defects in endosomal pH, LRP1 trafficking and amyloid clearance. Thus, NHE6 is a prominent effector of ApoE4 and emerges as a promising therapeutic target in Alzheimer disease.

## Introduction

Alzheimer disease (AD) is a degenerative brain disorder and a leading cause of dementia that affects 47 million people worldwide ^1^. AD is caused by pathological increase of amyloid β (Aβ) in the brain, resulting from an imbalance between its production and clearance. Recent studies suggest that accumulation of Aβ in the brain begins at least 20 years before symptoms appear ^2^. Although several promising drugs targeting the amyloid cascade have been developed, their astoundingly high failure rates (99.6%) in the clinic suggest that by the time amyloid plaques, neurofibrillary tangles and neuronal death are detected, it is unlikely that disease progression can be halted and reversed ^3^. Identifying and targeting pre-clinical pathologies may be critical for an effective cure.

In this context, endosomal aberrations constitute the earliest detectable brain cytopathology, emerging before cognitive dysfunction is apparent in neurodegenerative disorders, including Alzheimer disease, Niemann-Pick Type C and Down syndrome ^4, 5, 6, 7^. Consistent with this finding, genes associated with endosomal trafficking have been implicated as major risk factors in AD ^8^. Importantly, prominent pre-symptomatic endosomopathy of the brain, evidenced by enlarged and more numerous endosomes, has been observed in the ApoE4 genotype ^9, 10, 11^, the strongest known genetic risk factor influencing susceptibility to sporadic, late-onset AD (LOAD) ^12, 13^. The pathological E4 variant of Apolipoprotein E is present in ~50% of patients with AD, and the presence of two copies of the E4 allele increases risk of LOAD by ~12-times as compared to E3 isoform ^14^. Yet, despite strong evidence implicating endosomal uptake and clearance of Aβ in mediating AD risk in the ApoE4 genotype ^14, 15, 16^, the underlying mechanism is unknown.

Here we show that a profound dysregulation of endosomal pH in humanized mouse ApoE4 astrocytes leads to intracellular sequestration and cell surface loss of the Aβ receptor, LRP1. Selective down regulation of the endosomal Na^+^/H^+^ exchanger NHE6, which functions as a safety valve against excessive acidification by exchanging lumenal protons with cations, is mediated by increased nuclear translocation of the histone deacetylase HDAC4 in ApoE4 astrocytes. HDAC inhibitors that restore NHE6 expression also restore surface expression of LRP1 and effectively correct defective amyloid clearance in ApoE4 astrocytes to non-pathological ApoE3 levels. Consistent with a role in amyloid pathology, *in vivo* Aβ levels were found to be significantly higher in the brains of NHE6^KO^ mice. These findings could have implications for Christianson syndrome patients who have loss of function mutations in NHE6 and exhibit age-dependent hallmarks of neurodegeneration^17, 18^. In summary, we identify NHE6 as a novel ApoE effector and suggest potential therapeutic options in the treatment of amyloid disorders.

## Results

### ApoE4 astrocytes have cargo specific defects in endocytosis

Studies in human, mouse models and cell-culture have revealed the importance of ApoE-isotype-specific differences in Aβ uptake and clearance in AD pathogenesis, although the underlying mechanism remains to be determined ^14, 16, 19, 20^. To this end, we developed a sensitive and quantitative fluorescent-based assay to monitor cell-associated Aβ peptide (Fig. 1A) in astrocytes from ApoE^KO^ mice with isogenic knock-in of human ApoE3 and ApoE4 variants ^14^. Internalized Aβ is sorted to the lysosomal degradation pathway as evidenced by high colocalization with late endosomal-lysosomal markers and low colocalization with the recycling compartment marker transferrin (TFN) (Fig. S1A-D). Strikingly, cell-associated Aβ was reduced by 78% in ApoE4 astrocytes, relative to ApoE3 (Fig. 1B). To distinguish between Aβ uptake and turnover, we monitored the time course of Aβ internalization by flow cytometry analysis (Fig. 1C) and confocal microscopy (Fig. S1E). Consistent with defective uptake, there was significantly lower cell-associated Aβ in ApoE4 cells relative to ApoE3 at all time points (Fig. 1C, Fig. S1F). In contrast, cell-associated TFN was 1.5-2 fold higher in ApoE4 cells relative to ApoE3 as measured by flow cytometry (Fig. 1D) and confocal microscopy (Fig. 1E). Uptake of dextran by fluid-phase endocytosis was not different between ApoE genotypes (Fig. 1F). These observations reveal cargo selective effects of ApoE isotype in astrocytes and point to alterations in specific receptor pathways.

**Fig. 1.**
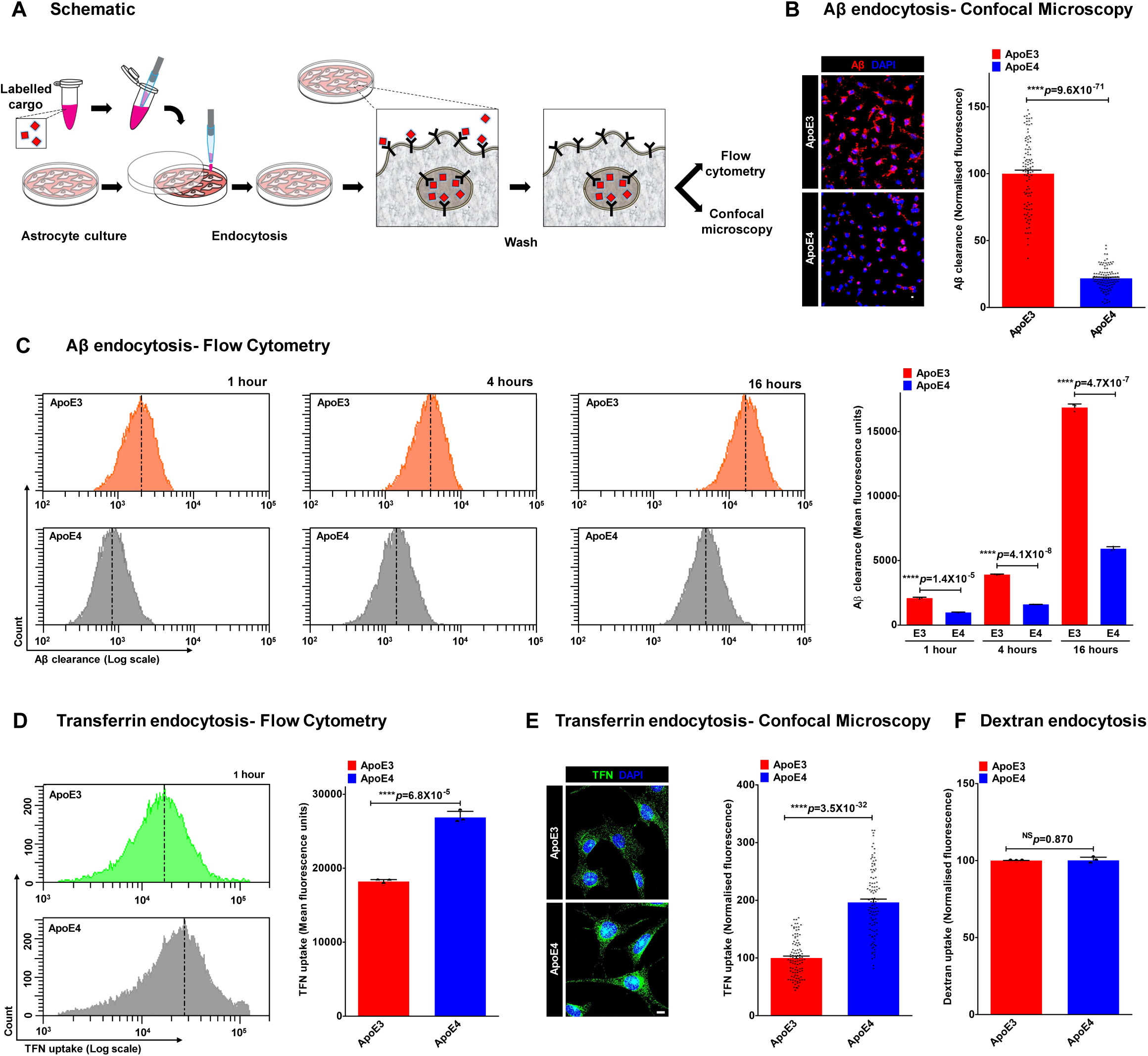
ApoE isotype-specific differences in Aβ clearance and specific receptor pathways. A. Fluorescent-based assay to monitor clearance of Aβ peptides by astrocytes. B. Representative micrographs (left) and quantification (right) of ApoE3 and ApoE4 astrocytes subjected to 24h of Aβ uptake. Fluorescence intensity and exposure settings were kept constant. Following background subtraction, fluorescence signal from cell associated Aβ was reduced by 78% in ApoE4 astrocytes, relative to ApoE3 (****p=9.6×10^−71^; *n*=100/condition; Student’s t-test). C. Representative fluorescence-activated cell sorting (FACS) histograms (left) demonstrating Aβ internalization by ApoE3 (top; orange) and ApoE4 astrocytes (bottom; grey) at 1h, 4h, and 16h; 10,000 cells/experimental condition. x-axis depicts Aβ clearance in logarithmic scale and vertical dashed line represents median fluorescence intensity. Quantification (right) of biological triplicate measurements of Aβ clearance from FACS analysis of ApoE3 and ApoE4 cells. Note significantly lower cell-associated Aβ in ApoE4 relative to ApoE3 at all time points (53% lower, 1 h; 59% lower, 4 h; 65% lower, 16 h; ****p<0.0001; *n*=3; Student’s t-test). D. Representative FACS histograms (left) and quantification of mean fluorescence intensity of biological triplicates (right) demonstrating TFN uptake following 60 minutes of endocytosis by ApoE3 (green) and ApoE4 (grey) astrocytes (~1.5-fold higher; ****p=6.8×10^−5^; *n*=3; Student’s t-test). x-axis of the FACS histogram depicts TFN uptake in logarithmic scale and vertical dashed line represents median fluorescence intensity. E. ApoE3 and ApoE4 astrocytes were incubated with fluorescent transferrin (TFN) for 1h, to compare steady-state TFN uptake by confocal microscopy. Fluorescence intensity and exposure settings were kept constant. Representative images are shown (left) and mean fluorescence ± s.e. was plotted (right). Following background subtraction, fluorescence signal was increased by ~2-fold in ApoE4 astrocytes, relative to ApoE3 (****p=3.5×10^−32^; *n*=100/condition; Student’s t-test). F. Quantification of mean fluorescence intensity of biological triplicates demonstrating dextran uptake by ApoE3 and ApoE4 astrocytes (p=0.870; *n*=3; Student’s t-test). See related Supplementary Fig. S1.

### Surface expression of LRP1 receptor is severely reduced in ApoE4 astrocytes

Transcriptional down regulation of the LRP1 receptor has been suggested as an underlying mechanism for defective Aβ clearance in AD patients^21^. However, we found no difference in brain LRP1 gene expression at different stages of AD (incipient, moderate, and severe), as compared with normal controls, in publicly available microarray data ^22^ (Fig. S2A-B). Meta-analysis of nine independent gene expression studies from anatomically and functionally distinct brain regions, comprising a total of 103 AD and 87 control post-mortem brains also showed no significant changes in LRP1 gene expression in AD (Fig. S2C-D). Consistent with these findings, we observed no differences in LRP1 transcript and total protein expression between ApoE3 and ApoE4 astrocytes (Fig. S2E-G).

LRP1 undergoes constitutive endocytosis from the membrane and recycling back to the cell surface ^23^. Therefore we considered the possibility that alterations in LRP1 receptor recycling could result in differences in plasma membrane expression. ApoE isotype-specific surface expression of LRP1 was evaluated using four independent approaches (Fig. 2A). First, surface biotinylation revealed that plasma membrane expression of LRP1 receptor in ApoE4 astrocytes was lower by ~50% (Fig. 1B). Second, an antibody directed against an external epitope of LRP1 to quantify surface expression in live cells by flow cytometry analysis showed a reduction of LRP1-positive cells by ~43% (Fig. 1C). Third, this was confirmed by confocal microscopy showing ~49% lower LRP1 surface labeling by antibody in ApoE4 (Fig. 1D). In a fourth approach, surface-bound ligand (fluorescent Aβ) measured by confocal microscopy was 66% lower in ApoE4 astrocytes (Fig. 1E). Notably, the greater attenuation in Aβ binding when compared to the ~50% reduction in surface LRP1 levels suggests additional isotype-specific mechanisms that contribute to Aβ clearance, such as reduced ligand-receptor affinity in ApoE4 cells or reductions in other Aβ receptors. Thus, ApoE isotype-specific alterations in receptor recycling determine LRP1 surface expression and cellular Aβ uptake, revealing a new pharmacological target for amyloid clearance defects in the pathological ApoE4 genotype.

**Fig. 2.**
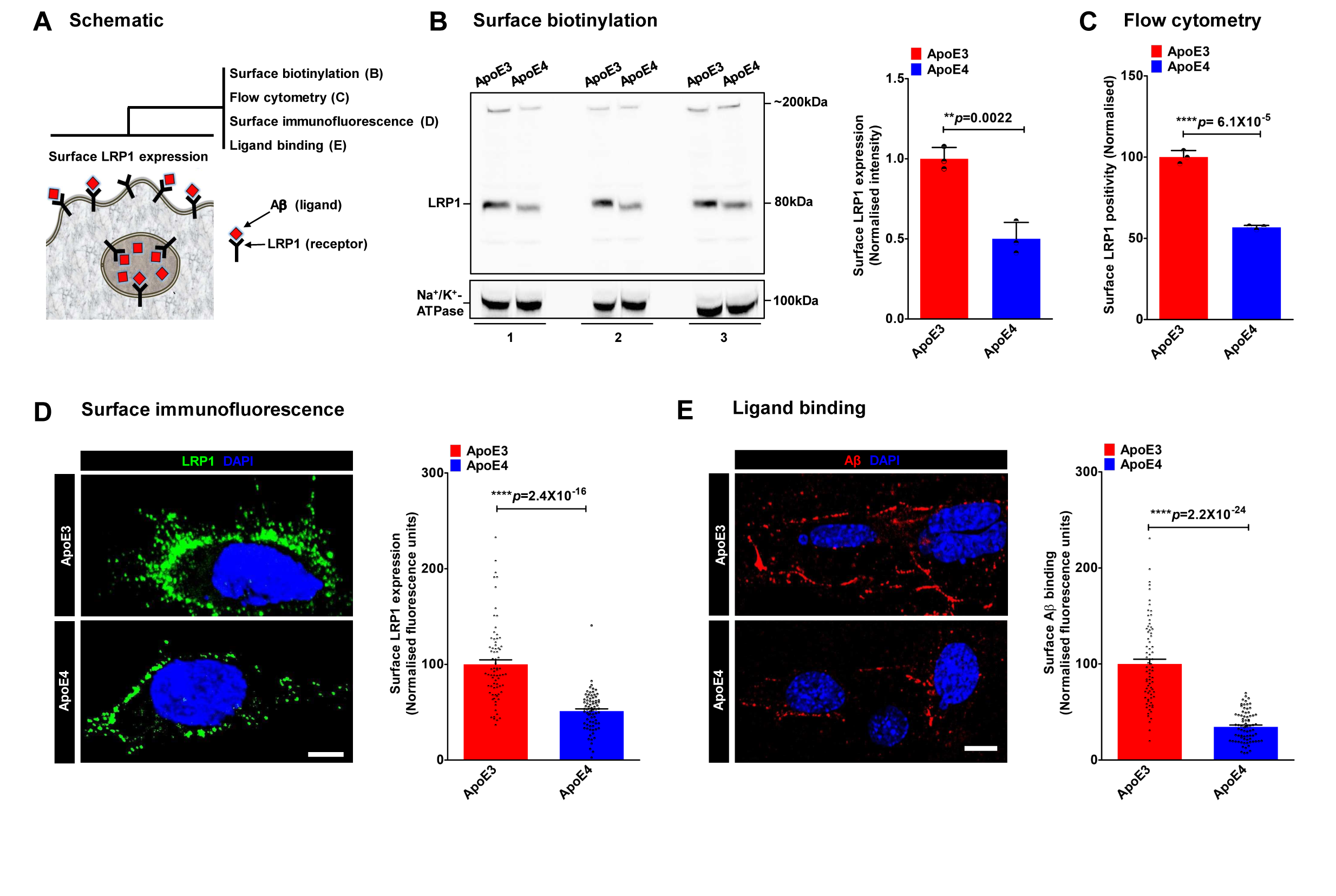
Reduced surface expression of LRP1 receptor in ApoE4 astrocytes. A. Four independent approaches to evaluate ApoE isotype-specific surface expression of LRP1. B. Surface biotinylation (left) and quantification (right) of biological triplicates showing that plasma membrane levels of LRP1 are depressed by ~50% in ApoE4 astrocytes, relative to ApoE3 (**p=0.0022; *n*=3; Student’s t-test). Plasma membrane protein Na^+^/K^+^ ATPase is used as loading control. C. Fraction of LRP1 positive cells quantified following surface antibody labeling and FACS analysis of 10,000 live, non-permeabilized cells in biological triplicates. Unstained cells were used as control. Note ~43% lower surface LRP1 positivity in ApoE4 relative to ApoE3 (****p=6.1×10^−5^; *n*=3; Student’s t-test). D. Representative surface immunofluorescence micrographs (left) and quantification (right) showing prominent LRP1 staining on cell surface and processes and faint, ~49% lower, labeling on ApoE4 cells (****p=2.4×10^−16^; *n*=75/condition; Student’s t-test). E. Plasma membrane level of LRP1 receptor was monitored by a ligand (fluorescent Aβ) binding assay performed on ice that only allows Aβ to bind surface receptors. Representative images are shown (left) and mean fluorescence ± s.d. was plotted (right). Surface-bound Aβ was 66% lower in ApoE4 astrocytes, relative to ApoE3 (*n*=75/condition; ****p=2.2×10^−24^; Student’s t-test). Scale bars, 10μm. See related Supplementary Fig. S2.

### Endo-lysosomal pH is defective in ApoE4 astrocytes

The pH within the endo-lysosomal system plays a critical role in receptor-mediated endocytosis and recycling ^24^. We used compartment-specific, pH-sensitive fluorescence reporters to probe ApoE-isotype dependent differences in endosomal, lysosomal and cytoplasmic pH (Fig. 3A). Endosomal pH in ApoE4 astrocytes was strongly reduced by ~0.84 pH unit, relative ApoE3 (Fig. 3B). In contrast, we observed >1 pH unit elevation of lysosomal pH in ApoE4 astrocytes (Fig. 3C). Previously, elevated lysosomal pH was observed in presenilin 1 (PS1)-deficient cell culture models and neurons, another genetic model of AD ^25^. Cytoplasmic pH showed no significant differences between the two ApoE isotypes (Fig. 3D).

**Fig. 3:**
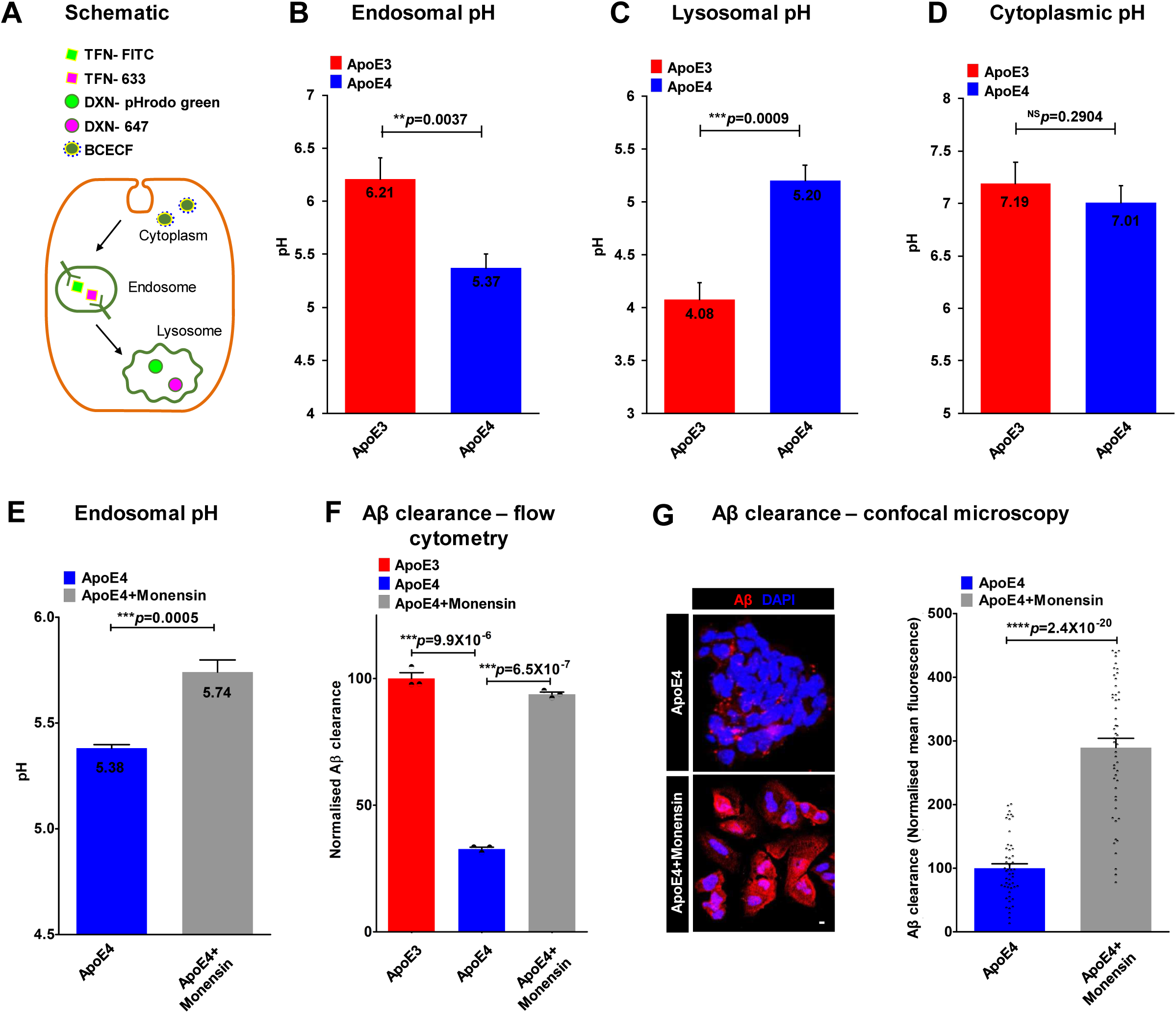
Endo-lysosomal pH is defective in ApoE4 astrocytes. A. Compartment-specific, ratiometric, pH-sensitive fluorescence reporters to probe ApoE-isotype dependent differences in endosomal, lysosomal and cytoplasmic pH. Endosomal pH was measured by incubations with pH-sensitive FITC-Transferrin (TFN-FITC) together with pH non-sensitive Alexafluor 633-Transferrin (TFN-633). Lysosomal pH was measured by incubations with pH-sensitive pHrodo-green-Dextran (DXN-pHrodo green) together with pH non-sensitive Alexa Fluor 647-Dextran (DXN-647). Cytoplasmic pH was measured ratiometrically using pH-sensitive green and pH nonsensitive red fluorescence of BCECF dye. B. Endosomal pH in ApoE4 astrocytes was strongly reduced by ~0.84 pH unit, relative to ApoE3 (**p=0.0037; *n=3*; Student’s t-test). C. Lysosomal pH was elevated by >1 pH unit in ApoE4 astrocytes (****p=0.0009; n=3; Student’s t-test). D. Cytoplasmic pH showed no significant differences between ApoE3 and ApoE4 astrocytes (p=0.2904; n=3; Student’s t-test). E. Monensin treatment (50μM for 1h) corrected hyperacidic endosomal pH in ApoE4 astrocytes, relative to vehicle treatment (****p=0.0005; n=3; Student’s t-test). F. Quantitation of Aβ clearance from FACS analysis of 10,000 cells in biological triplicates confirmed restoration of Aβ clearance in ApoE4 astrocytes to ApoE3 levels with monensin treatment (****p=6.5×10^−7^; *n*=3; Student’s t-test). G. Representative micrographs (left) and quantification (right) showing ~2.9-fold increase in cell-associated Aβ in ApoE4 astrocytes with monensin treatment (****p=2.4×10^−20^; *n*=50; Student’s t-test). Scale bars, 10μm.

To determine if there was a causal link between endo-lysosomal pH and defective Aβ clearance in ApoE4 astrocytes, we treated ApoE4 cells with the ionophore monensin that mediates Na^+^/H^+^ exchange across acidic compartments^26^. Thus, monensin treatment (50μM for 1 h) elevated endosomal pH in ApoE4 knock-in astrocytes from 5.38±0.01 to 5.74±0.03, relative to the vehicle treated control (Fig. 3E). Concomitantly, monensin treatment restored Aβ clearance in ApoE4 astrocytes to ApoE3 levels, as shown by flow cytometry analysis (Fig. 3F). This was independently confirmed by confocal microscopy (Fig. 3G), suggesting that defective pH regulation could underlie the observed Aβ clearance defects.

### NHE6 restores defective Aβ clearance in ApoE4 astrocytes

Luminal pH in the endo-lysosomal network is set by the precise balance of proton pump and leak pathways, mediated by V-type H^+^-ATPase and endosomal Na^+^/H^+^ exchangers (NHE6), respectively (Fig. 4A)^27, 28, 29, 30^. Changes in expression and activity of the pump and leak pathways could lead to significant dysregulation of endosomal pH in Alzheimer brains. Consistent with this possibility, analysis of a publicly available microarray dataset (GSE5281) comprising a total of 15 sporadic, late-onset AD (LOAD) and 12 matched control post-mortem brains ^31^ revealed that genes involved in hydrogen ion transmembrane transport, including the endosomal NHE6 and V-ATPase subunits, comprised 10% of the top 100 down regulated genes, exhibiting highest enrichment scores (>15-fold; Fig. S3A). In AD patients with ApoE4/4 genotype, NHE6 was among the transcripts differentially down regulated in hippocampus, by up to ~4-fold compared to ApoE3/3 ^32, 33^. We validated these findings using an independent, large human brain transcriptome dataset (*n*=363, GSE15222) to show ApoE4 isotype-specific differential gene expression of NHE6 in aging brain ^34^ (Fig. 4B). Although NHE6 transcript was similar in ApoE^KO^ mouse astrocytes and ApoE^KO^ astrocytes with knock-in of human ApoE3, it was ~56% reduced in ApoE4 knock-in cells (Fig. 4C). There was also ApoE4 specific reduction in transcript for the related endosomal isoform NHE9 (~70% lower) and lysosomal V-ATPase V0a1 subunit (~67%), but not for the plasma membrane NHE1 isoform (Fig. S3B). These large transcript differences could account for the observed ApoE-isotype specific shifts in endo-lysosomal pH.

**Fig. 4:**
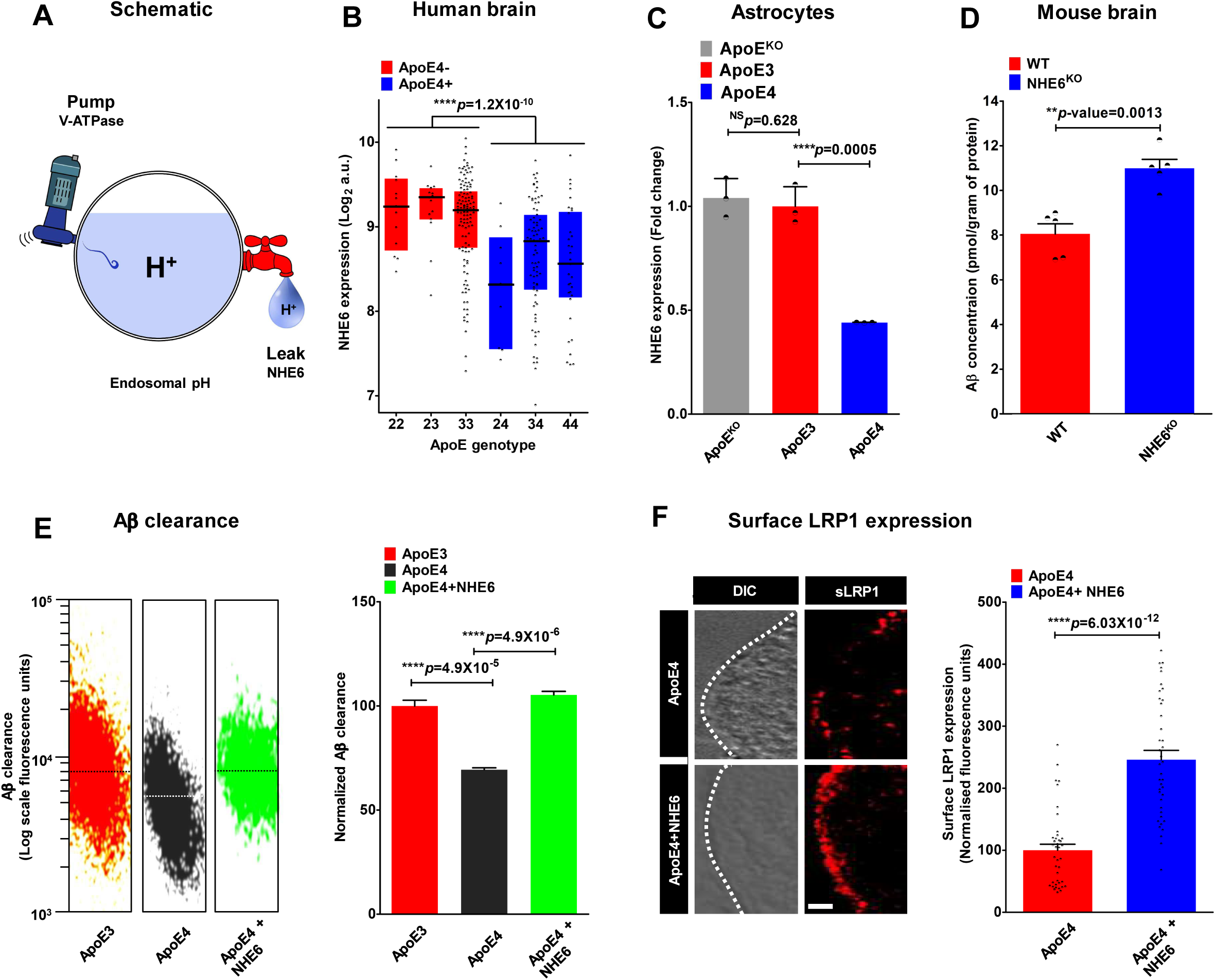
NHE6 restores defective Aβ clearance in ApoE4 astrocytes. A. Endosomal pH is precisely tuned by a balance of proton pumping (acidification) through the V-ATPase and proton leak (alkalization) via NHE6. B. Box-plots of NHE6 transcript levels in postmortem brains extracted from a large microarray dataset (*n*=363, GSE15222) showing significant downregulation of NHE6 expression in ApoE4+ brains (****p=1.2×10^−10^; Student’s t-test). C. Quantitative PCR (qPCR) analysis of NHE6 transcript reveled significantly lower expression in ApoE4 astrocytes, relative to ApoE3 (~56% lower, ****p=0.0005, n=3; Student’s t-test). No difference in NHE6 levels was observed between ApoE3 astrocytes and isogenic ApoE^KO^ astrocytes (p=0.628; *n*=3; Student’s t-test). D. Aβ levels were significantly higher in the brains of NHE6-null mouse model (NHE6^KO^), relative to WT (**p=0.0013; *n*=5/condition; Student’s t-test), consistent with our hypothesis. Aβ was measured using ELISA and normalized to total protein concentration in the brain homogenate. E. FACS scatter plots (left) and quantification (right) demonstrating Aβ internalization by empty vector expressing ApoE3 (orange) and ApoE4 (grey) astrocytes and ApoE4 astrocytes with restored NHE6 expression (green). Note remarkable correction of Aβ clearance in ApoE4 astrocytes with NHE6 expression to ApoE3 levels (****p=4.9×10^−6^; *n*=3; Student’s t-test). F. Representative surface immunofluorescence images (left) and quantification (right) of non-permeabilized cells showing ~2.5-fold increase in plasma membrane LRP1 expression in ApoE4 cells expressing exogenous NHE6 (****p=6.02×10^−12^; *n*=40/condition; Student’s t-test). Scale bars, 2.5μm. See related Supplementary Fig. S3.

Taken together, these data suggest an important, hitherto underappreciated role of proton transport and endosomal pH regulation in AD. We hypothesized that NHE6 is a potential ApoE effector, and that down regulation of NHE6 in disease-associated ApoE4 variants is causal to a subset of AD phenotypes. Consistent with this hypothesis, amyloid Aβ levels were found to be elevated in mouse brains from NHE6^KO^ (Fig. 4C), together with diminished brain weight (Fig. S3C), suggesting an underlying neurodegenerative pathology.

Similar to monensin treatment, lentiviral vector mediated expression of NHE6 alkalinized the endosomal lumen in ApoE4 astrocytes (Fig. S3D). Therefore, we tested if ectopic expression of GFP-tagged eNHE isoforms could correct defective Aβ uptake in ApoE4 astrocytes. Remarkably, Aβ clearance was restored to ApoE3 levels in ApoE4 astrocytes transfected with NHE6 (Fig. 4E), but not with NHE9 (Fig. S3E-F), pointing to an isoform-specific role for NHE6 in Aβ clearance.

Colocalization of NHE6 with EEA1 and LRP1 (Fig. S3G-H) suggested a potential role for NHE6 in endosomal recycling of LRP1 receptors. Compared to the weak surface LRP1 staining in vector-transfected ApoE4 astrocytes, we observed prominent, ~2.5-fold higher LRP1 staining in ApoE4 cells expressing ectopic NHE6 (Fig. 4F). Similar results were obtained in surface biotinylation experiments that showed robust ~5.7-fold higher surface LRP1 levels in ApoE4 cells with restored NHE6 expression, compared to transfection with empty vector (Fig. S3I). We confirmed that there were no concomitant changes in LRP1 transcript (Fig. S3J) or total protein expression levels (Fig. S3I), suggesting that increased surface LRP1 was due to posttranslational redistribution of the existing cellular LRP1 pool. Taken together, our data point to diminished NHE6 expression as a major underlying cause for defective Aβ clearance in ApoE4 astrocytes. Furthermore, since LRP1 is a receptor for multiple ligands, loss of NHE6 may contribute to other ApoE4 defects, including defective synaptosome uptake and synapse pruning ^12, 23^.

### HDAC inhibitors rescue NHE6-mediated Aβ clearance deficits

Reports of increased nuclear translocation of multiple histone deacetylases (HDACs) in ApoE4 isotype, relative to ApoE3 (Fig. 5A), in post-mortem brains and neurons suggested a mechanistic basis for our observations ^35^. Fractional colocalization of HDAC4 with DAPI revealed prominent overlap, consistent with increased nuclear translocation in ApoE4 astrocytes (Fig. 5B). This was independently verified in Western blots of nuclear fractions, which showed higher HDAC4 in ApoE4 astrocytes relative to ApoE3 (Fig. S4A).

**Fig. 5:**
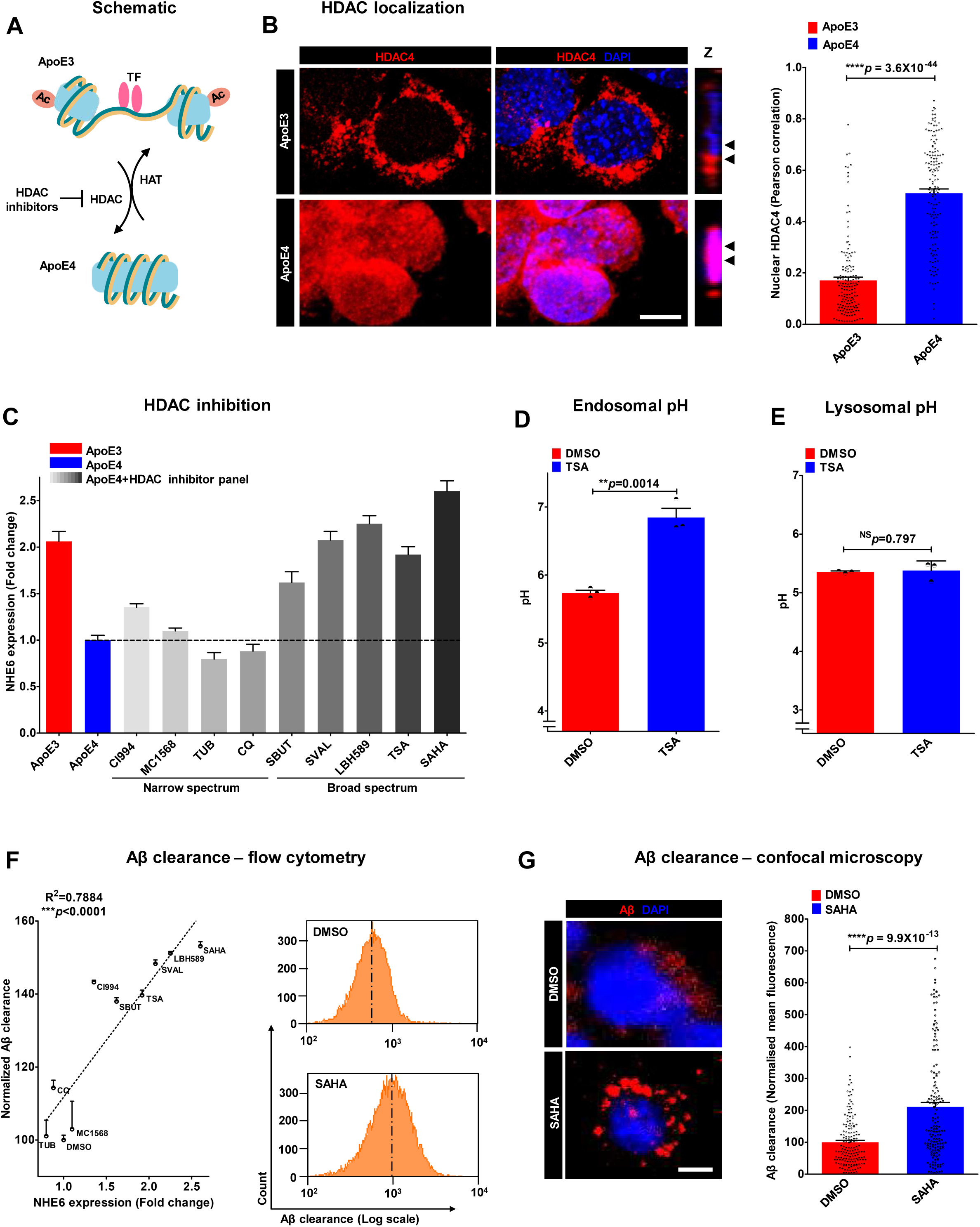
HDAC inhibitors rescue NHE6-mediated Aβ clearance deficits. A. ApoE4 increases nuclear translocation of histone deacetylases (HDACs), whereas ApoE3 increases histone acetylation and transcription factor (TF) binding. Therapeutic effects of HDAC inhibitors are potentially mediated by reducing effects of ApoE4-induced nuclear translocation of HDACs. B. Representative micrographs (left) and quantification using Pearson correlation coefficient (right) determining fractional colocalization of HDAC4 (red) with DAPI (blue) in ApoE3 and ApoE4 astrocytes. Colocalization is evident in the merge and orthogonal slices (Z) as magenta puncta. Note prominent overlap between HDAC4 and DAPI, consistent with increased nuclear translocation, in ApoE4 astrocytes (Pearson’s correlation coefficient= ApoE3: 0.17±0.01 vs. ApoE4: 0.51±0.02; n=150/condition; ****p=4.6×10^−44^; Student’s t-test). C. qPCR analysis to determine the potential of HDAC inhibitor panel to augment the expression of NHE6 in ApoE4 astrocytes following 12h treatment. Note that inhibitors of class I (CI994) or class II (MC1568) HDACs resulted in minimal changes in NHE6 expression. Broad-spectrum drugs inhibiting both classes, including sodium butyrate, sodium valproate, LBH589, TSA, and SAHA, resulted in significant restoration of NHE6 expression levels in ApoE4 astrocytes to levels comparable to ApoE3 astrocytes (****p<0.0001; *n*=3; Student’s t-test). Other narrow spectrum HDAC inhibitors studied here (tubacin/TUB and clioquinol) had no significant effect. D. HDAC inhibition by TSA treatment (5μM for 12h) corrected hyperacidic endosomal pH in ApoE4 astrocyte, relative to DMSO treatment (**p=0.0014; *n*=3; Student’s t-test). E. TSA treatment (5μM for 12h) did not affect lysosomal pH in ApoE4 astrocyte, relative to DMSO treatment (p=0.797; *n*=3; Student’s t-test). F. Quantitation of Aβ clearance from FACS analysis of 10,000 cells in biological triplicates to determine efficacy of panel of HDAC inhibitors to rescue Aβ clearance deficits in ApoE4 astrocytes. Note prominent linear relationship between Aβ clearance and the fold-change in NHE6 expression (R^2^=0.7884; p<0.0001) elicited by DMSO and nine HDAC inhibitors. Representative FACS histogram demonstrating increase Aβ internalization by ApoE4 with SAHA treatment is shown on the right. G. Representative micrographs (left) and quantification (right) demonstrating internalized Aβ following SAHA treatment. Note prominent, vesicular Aβ staining in SAHA treated ApoE4 cells, in contrast to faint, diffuse staining in vehicle control (****p=9.9×10^−13^; *n*=160/condition; Student’s t-test). Scale bars, 10μm. See related Supplementary Fig. S4.

To translate these observations, we screened a panel of nine HDAC inhibitors comprising several different chemical classes for their potential to augment the expression of NHE6 in ApoE4 astrocytes. Whereas inhibitors of class I (CI994) or class II (MC1568) HDACs resulted in minimal changes in NHE6 expression, broad-spectrum drugs inhibiting both classes, including sodium butyrate, sodium valproate, LBH589, TSA, and SAHA (vorinostat), resulted in significant restoration of NHE6 expression levels in ApoE4 astrocytes to levels comparable to ApoE3 astrocytes (Fig. 5C). Other narrow spectrum HDAC inhibitors studied here (tubacin and clioquinol) had no significant effect. Both TSA and SAHA elicited dose dependent NHE6 increases with half-maximal response (EC_50_) of 6.50±0.36μM and 6.81±0.53μM (Fig. S4B-C), respectively, comparable to their therapeutic plasma concentrations ^36^. Neither TSA nor SAHA significantly altered NHE9 levels (Fig. S4D). We confirmed that both TSA and SAHA stimulated acetylation of histone H3 and H4 in ApoE4 astrocytes following 60 minute of treatment (Fig. S4E-F). Next, we sought to determine if enhanced NHE6 expression resulting from inhibition of histone deacetylases was physiologically effective in correcting hyperacidic endosomal pH in ApoE4 astrocytes. TSA treatment (5μM for 12h) exhibited a compartment-specific effect of significantly elevating endosomal pH (Fig. 5D) without effect on lysosomal pH (Fig. 5E). Of note, TSA or SAHA treatment in ApoE4 astrocytes did not significantly affect cell viability measured using trypan blue exclusion (Fig. S4G).

Key to the potential efficacy of HDAC inhibitors in AD therapy is their ability to rescue Aβ clearance deficits in ApoE4 astrocytes. We observed a prominent linear relationship between Aβ clearance and the fold-change in NHE6 expression (R^2^=0.7884; Fig. 5F) elicited by the panel of nine HDAC inhibitors. HDAC inhibitors with lower induction of NHE6 expression (e.g. MC1568 and tubacin) conferred minimal changes in Aβ clearance. Notably, broad-spectrum HDAC inhibitors (e.g. TSA and SAHA) that significantly restored NHE6 expression also elicited proportionally complete correction of defective Aβ clearance in ApoE4 astrocytes to levels similar (up to 92.4%) to ApoE3 cells (Fig. 5F and S4H). ApoE4 cells treated with SAHA showed prominent, vesicular Aβ staining relative to vehicle control (Fig. 5G). Taken together, these findings reveal differential effects of ApoE3 and ApoE4 genotypes on nucleo-cytoplasmic shuttling of the HDACs leading to a novel molecular mechanism, with clinical implication, for ApoE4 associated down regulation of NHE6 in post-mortem brain and astrocyte models.

## Discussion

The discovery of endosomal Na^+^/H^+^ exchangers (eNHE) first in yeast, and soon after in plants, metazoans, and mammalian systems, established their evolutionarily conserved role as a leak pathway for protons in compartmental pH homeostasis, critical for cargo trafficking and vesicular transport ^27, 37, 38^. Na^+^/H^+^ exchangers are estimated to have exceptionally high transport rates of ~1,500 ions/s ^39^, so that even small perturbations in expression result in dramatic changes in ionic milieu within the limited confines of the endosomal lumen. Genetic studies have linked eNHE to a host of neurodevelopmental and neurodegenerative disorders, including Christianson syndrome (CS) with symptoms of autism, intellectual disability and epilepsy, Parkinsons disease, multiple sclerosis and AD, although underlying mechanisms remain to be determined ^27^.

One clue emerged from network analysis of 1697 genes in a late-onset AD dataset (Cases n=176, Controls=188), in which the endosomal Na^+^/H^+^ exchanger NHE6 *(SLC9A6)* was identified as a top-five hub transcript in AD, with 202 network connections and a plethora of potential downstream effects ^34^. More recent network analysis of the metastable subproteome associated with AD also converged on NHE6 as a major hub gene regulating protein trafficking and clearance mechanisms ^40^. In addition to prominent neurodegeneration phenotypes in CS patients, female carriers have learning difficulties and behavioral issues, and some present with low Mini Mental Status Exam (MMSE) scores suggestive of early cognitive decline ^41^. Interestingly, NHE6 was among the most highly down regulated genes (up to 6-fold) in elderly (70 years) brain, compared to adult (40 years)^42^. These observations point to a more widespread role for NHE6 in neurodegenerative disorders and unexpectedly common pathological pathways between CS and Alzheimer disease.

Previously, we showed that NHE6 regulates trafficking and BACE1-mediated processing of amyloid precursor protein APP to limit production of amyloidogenic peptides ^43^. In this study, we demonstrate a critical role for NHE6 in the uptake and clearance of soluble, secreted Aβ in astrocytes. Using ApoE-isoform-expressing isogenic astrocytes that produce, lipidate, package, and secrete ApoE in a brain-relevant physiological fashion, we showed that the well-documented pathogenic deficiency of ApoE4 astrocytes to clear Aβ is mediated by decreased expression of NHE6, which results in endosomal over-acidification and reduced surface levels of the Aβ receptor LRP1. Thus, NHE6 is a novel and important ApoE4 effector in astrocytes (Fig. 6). The precise pH-mediated perturbation in trafficking remains to be determined. It is likely that hyper-acidification of early and recycling endosomes redistributes plasma membrane proteins to the lysosome at the expense of recycling to the cell surface. Taken together, we propose that loss of NHE6 function contributes to the endosomal pathology observed in pre-symptomatic AD brains both by accelerating Aβ production and by inhibiting Aβ clearance, promoting the development of amyloid plaques and culminating in neurodegeneration and dementia.

**Fig. 6:**
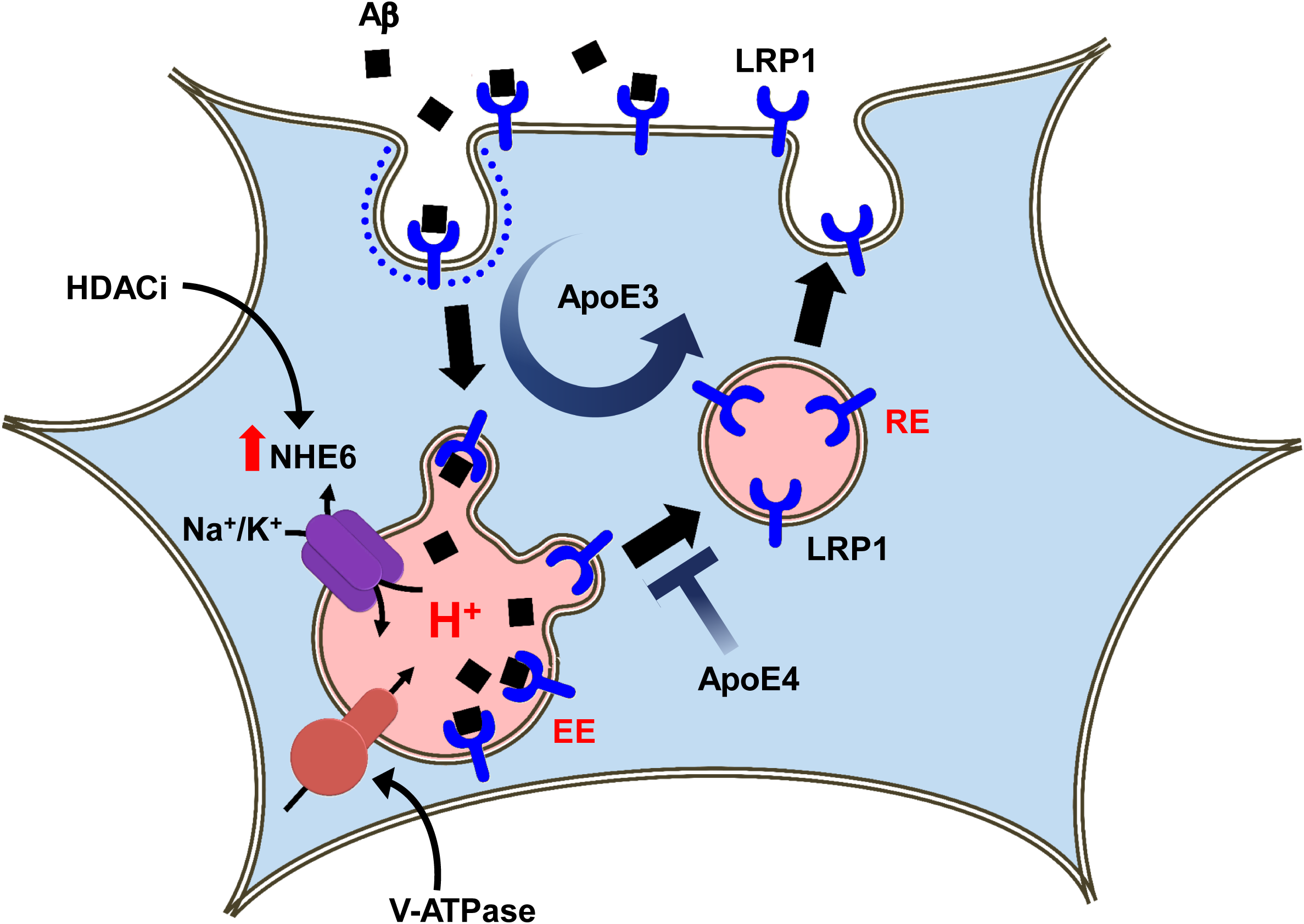
Proposed Role for NHE6 in Aβ clearance in astrocytes. Amyloid beta (Aβ) receptor LRP1 is constitutively recycled to the cell surface through early and recycling endosomes (EE and RE) in ApoE3 astrocytes. Loss of NHE6 expression in ApoE4 astrocytes hyperacidifies endosomes and impairs trafficking of LRP1 receptor resulting in defective Aβ clearance. Histone deacetylase inhibitors (HDACi) restore expression of NHE6 and Aβ clearance in ApoE4 cells.

Abnormalities in histone acetylation have been linked to several neurodegenerative diseases including AD, and HDAC inhibitors appear to show a neuroprotective effect, improving memory and cognition in mouse models ^44, 45^. Here, we link increased nuclear translocation of HDAC4 in ApoE4 astrocytes to down regulation of NHE6 expression. We show that broad-spectrum HDAC inhibitors restore NHE6 expression, normalize endosomal pH and correct Aβ clearance defects in ApoE4 astrocytes. Thus, the amelioration of AD pathogenesis observed *in vitro* and *in vivo* by small molecule inhibitors of HDACs may be mediated, in part, by NHE6. Future work could test the efficacy of these pharmacological agents on amyloid pathology in well-defined animal models. Given the well-known link between NHE6 dysfunction and epilepsy ^27, 46^, we suggest that increased NHE6 expression could potentially contribute to anti-epileptic mechanisms of HDAC inhibitor drug sodium valproate. Importantly, our data demonstrate a hitherto unrecognized ability of HDAC inhibitors to specifically enhance endosomal pH that could potentially correct human pathologies resulting from aberrant endosomal hyperacidification.

Dysfunction in endo-lysosomal pH is an emerging theme in AD with clear potential for intervention to exploit the disease-modifying effects of endosomal pH ^7^. Amphipathic drugs such as bepridil and amiodarone partition into acidic compartments, alkalinize endosomes, and correct Αβ pathology in cell culture and animal models ^47^. Our study supports a rational, mechanistic basis for such repurposing of existing FDA-approved drugs with well-established safety and pharmacokinetic profiles, known to have off-label activity of endosomal alkalization, to target the cellular microenvironment in AD. Similar to our observations in AD, down regulation of NHE6 gene expression has been reported in autism brains ^48^. We suggest that endosomal pH may be a critical mechanistic link between neurodevelopmental and neurodegenerative disorders. Thus, a subset of autism patients with dysregulated NHE6 activity, either from loss-of-function mutations or by down-regulated gene expression, are likely to have a high risk of developing neurodegenerative disorders, thereby providing a rational basis to stratify patients for targeted therapies. In conclusion, this work presents, (i) a novel ApoE4 regulated cellular mechanism and druggable target in AD, namely, regulation of amyloid pathology by intra-endosomal pH; (ii) a new focus on endosome trafficking in astrocyte function, a neglected area in neurodegenerative disorders; (iii) a new link between an autism gene and the AD risk allele ApoE4; (iv) a new strategy for mechanism-based therapies for AD and related devastating disorders with important implications for early intervention to limit progressive, severe and debilitating neurodegeneration seen in Christianson syndrome patients.

## Author Contributions

H.P. designed, conducted and analyzed experiments and wrote the paper. R.R. designed and interpreted experiments and wrote the paper.

## Acknowledgements

We thank Drs. Robert Edwards and Julie Ullman of University of California San Francisco for providing mouse brains and Dr. David M Holtzman, Washington University, St. Louis for the gift of ApoE immortalized astrocytes. We are very grateful to Dr. Seth S. Margolis for helpful discussions, and Richard L. Blosser for assistance with the flow cytometry analysis. This work was made possible by support from the Johns Hopkins Medicine Discovery Fund to R.R. Additional support came from a grant to R.R. from the National Institutes of Health (DK054214). H.P. is Fulbright Fellow supported by the International Fulbright Science and Technology Award.

---

**Fig. S1:**
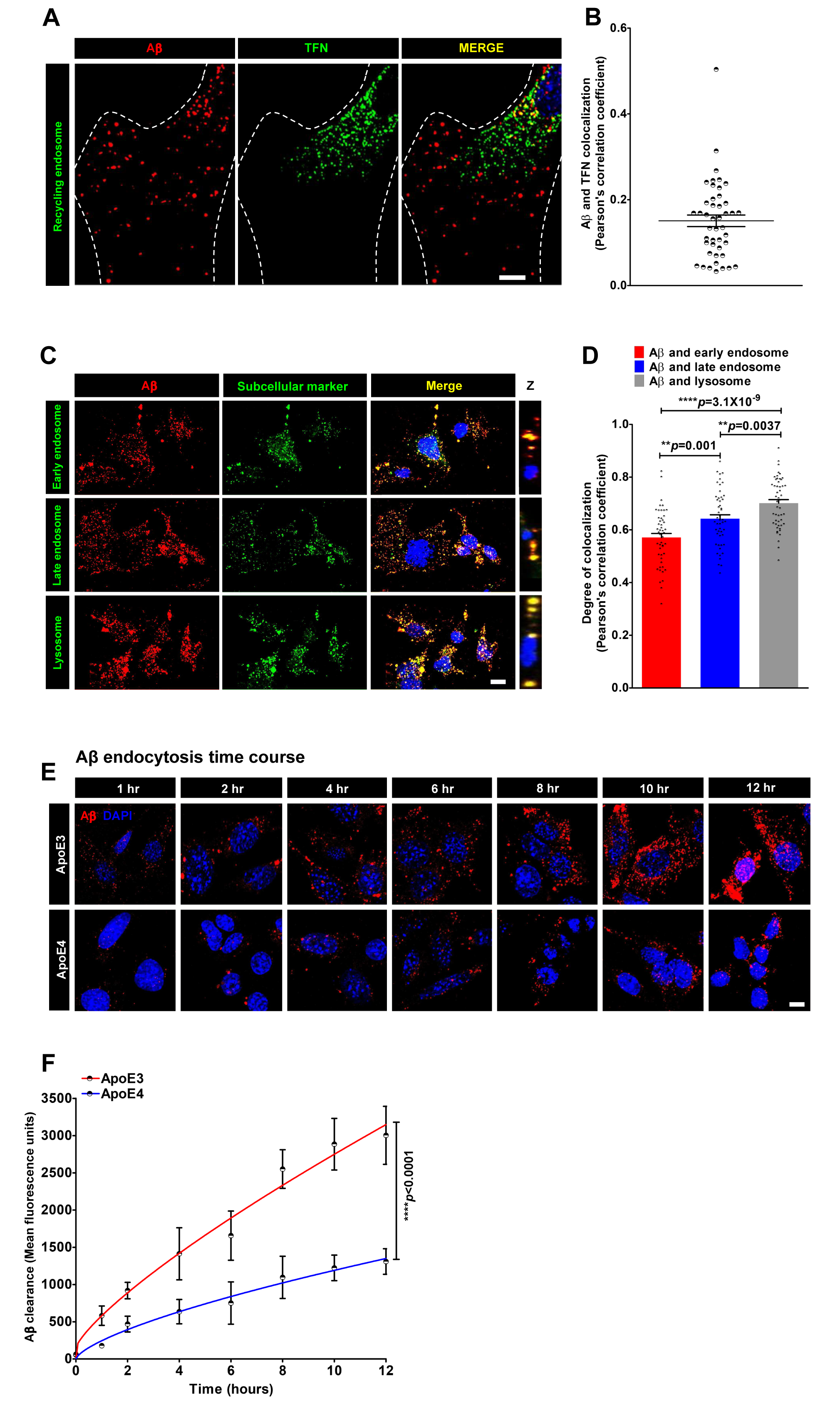
ApoE isotype-specific differences in Aβ clearance. Related to Fig 1. (A-B) Representative micrographs (A) and quantification using Pearson correlation (B) determining fractional colocalization of Aβ (red) with transferrin (TFN) (green) in DAPI-(blue) stained ApoE3 astrocytes, following 60 minutes of uptake. Note poor colocalization, as evident in the merge and orthogonal slices (Z) with fewer yellow puncta (Pearson’s correlation: 0.16±0.09; *n*=45), suggesting that a significant pool of internalized Aβ escapes recycling endosomes. Representative micrographs (C) and quantification using Pearson correlation (D) determining fractional colocalization of Aβ (red) with different endosomal-lysosomal compartment markers (green) in DAPI-(blue) stained ApoE3 astrocytes following 12h of uptake. Colocalization is evident in the merge and orthogonal slices (Z) as yellow puncta. (E-F) Time course of Aβ accumulation in ApoE3 and ApoE4 astrocytes by confocal microscopy. Representative images are shown (E) and mean fluorescence ± s.e. was plotted (F). ApoE4 cells show a reduced rate and total Aβ accumulation, relative to ApoE3 astrocytes (****p<0.0001; Student’s t-test). Scale bars, 10μm.

**Fig. S2:**
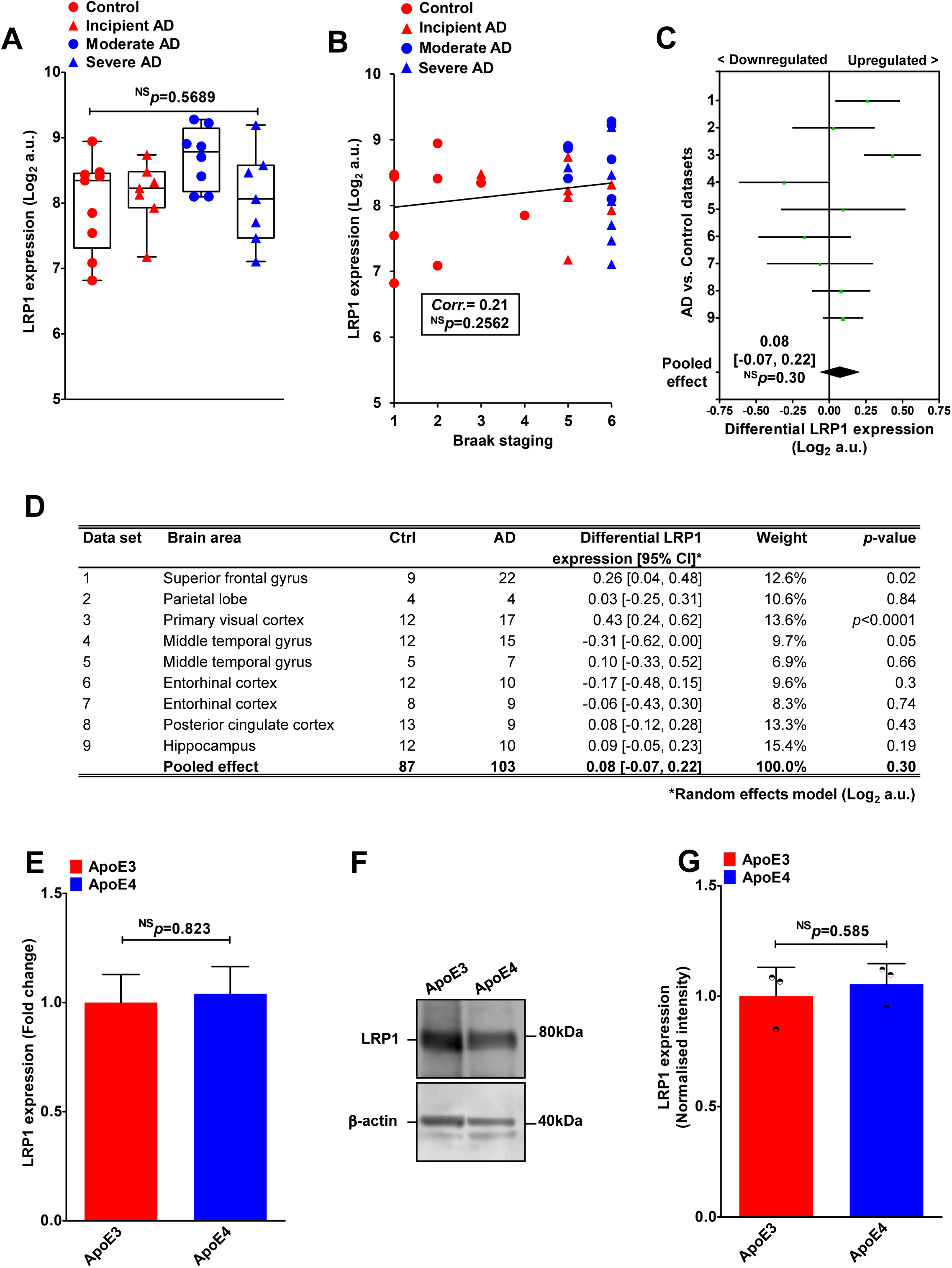
LRP1 transcript and total protein levels in ApoE isotypes. Related to Fig 2. A. Box and whisker plots of hippocampal LRP1 gene expression across control and clinical stages of (p=0.569; *n*=31; ANOVA). B. Expression plot of LRP1 with post-mortem brain pathology assessed via Braak staging scores across all 31 subjects, regardless of diagnosis. Scores on the Braak staging range from 0 to 6, with higher scores indicate worse neuropathology. No correlation was observed for LRP1 levels with Braak stage (Pearson correlation coefficient=0.21; *n*=31; p=0.2562). Black line depicts linear fit. (C-D) Forest plot (C) and quantification (D) of differential LRP1 expression between AD and control (Ctrl) data sets and pooled average represented as mean difference and 95% confidence interval (CI) of log base 2 expression, were obtained as described under “Experimental Methods.” (Black diamond/pooled average=0.08; 95% CI= −0.07, 0.22; control, *n*=87; AD patients, *n*=103; p=0.30). E. LRP1 gene expression showing no significant difference between ApoE3 and ApoE4 astrocytes (p=0.823; Student’s t-test). Representative Western blot (F) and quantification of three biological replicates (G) showing total LRP1 protein expression normalized to β-actin levels between ApoE3 and ApoE4 astrocytes (p=0.585; *n*=3; Student’s t-test).

**Fig. S3:**
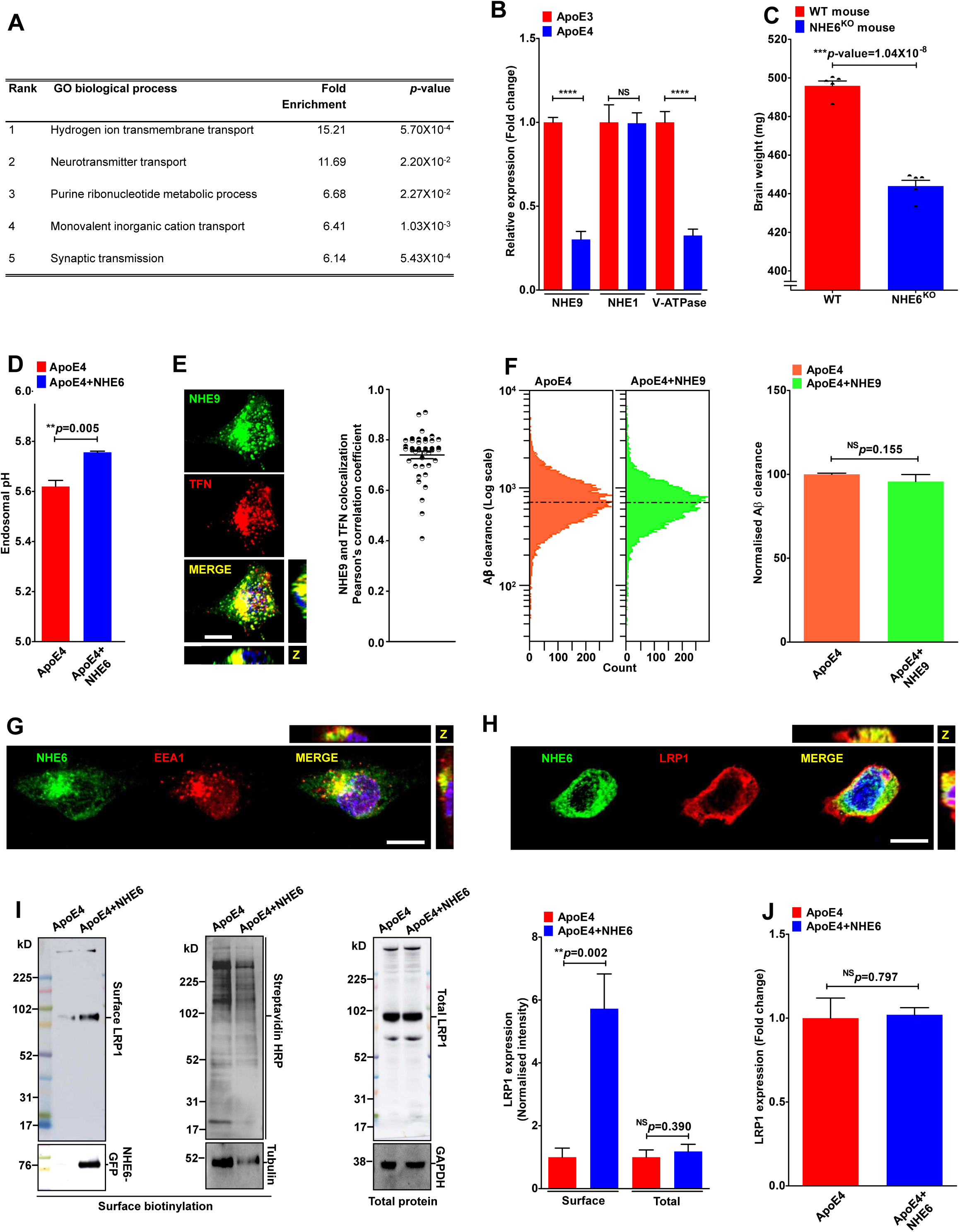
NHE6 corrects defective surface LRP1 expression in ApoE4 astrocytes. Related to Fig 4. A. Gene ontology (GO) analysis of top-100 downregulated genes in post-mortem AD brains obtained as described under “Experimental Methods.” Enrichment scores and p-values for top-5 GO biological process were shown. Note that genes involved in hydrogen ion transmembrane transport, including NHE6 and V-ATPase subunits exhibited highest enrichment scores. B. Quantitative PCR (qPCR) analysis of NHE9 transcript reveled significantly lower expression in ApoE4 astrocytes, relative to ApoE3 (~70% lower; ****p=2.7×10^−5^; *n*=3; Student’s t-test). No changes in mRNA levels were observed for the closely related plasma membrane NHE1 isoform (p=0.946; *n*=3; Student’s t-test). Significantly lower mRNA levels for lysosomal V-ATPase V0a1 subunit in ApoE4 astrocytes, relative to ApoE3 (~67% lower; ****p=9.7×10^−5^; *n*=3; Student’s t-test). C. Consistent with clinical reports of microcephaly in Christianson syndrome patients, 7-month old hemizygous NHE6^KO^ mice showed significantly lower brain weight, relative to wild-type mice. D. Lentiviral vector mediated expression of NHE6 in ApoE4 astrocytes with low endogenous NHE6 levels results in alkalization of endosomal pH. E. Representative micrographs (left) and quantification using Pearson correlation (right) determining fractional colocalization of NHE9 (green) with transferrin (TFN) (red) in DAPI-(blue) stained ApoE4 astrocytes following 60 minutes of uptake. Note prominent endosomal colocalization of NHE9 as evident in the merge and orthogonal slices (Z) as yellow puncta (Pearson’s correlation: 0.74±0.10; *n*=40). F. Representative FACS histograms (left) and quantification of mean fluorescence intensity of biological triplicates (right) demonstrating no difference in Aβ internalization between ApoE4 (red) and ApoE4 astrocytes with NHE9-GFP expression (green). x-axis depicts Aβ clearance in logarithmic scale and vertical dashed line represents median fluorescence intensity (p=0.155; *n*=3; Student’s t-test). (G-H) Representative immunofluorescence images of permeabilized, fixed ApoE4 astrocytes expressing NHE6-GFP showing overlap of NHE6 (green) with red-labeled EEA1 (G) and LRP1 (H). Colocalization is evident in the merge and orthogonal slices (Z) as yellow puncta. I. Representative blots showing surface (left) and total (right) LRP1 protein levels with NHE6-GFP expression (detected using anti-GFP antibody) relative to empty vector transfection. As loading control, surface biotinylated proteins were visualized with HRP-streptavidin and by probing against tubulin. GAPDH was used as a loading control for western analysis of total LRP1 levels. Quantification (extreme right) of blots showed robust ~5.7-fold higher surface LRP1 levels in ApoE4 cells with restored NHE6 expression, compared to transfection with empty vector (**p=0.002; *n*=3; Student’s t-test). No concomitant changes in total LRP1 levels (p=0.390; *n*=3; Student’s t-test) suggesting that increased surface LRP1 was due to posttranslational redistribution of the existing cellular LRP1 pool. J. qPCR analysis confirmed no concomitant changes in LRP1 transcript with lentiviral transduction of NHE6-GFP, relative to empty vector control (p=0.797; *n*=3; Student’s t-test). Scale bars, 10μm.

**Fig. S4:**
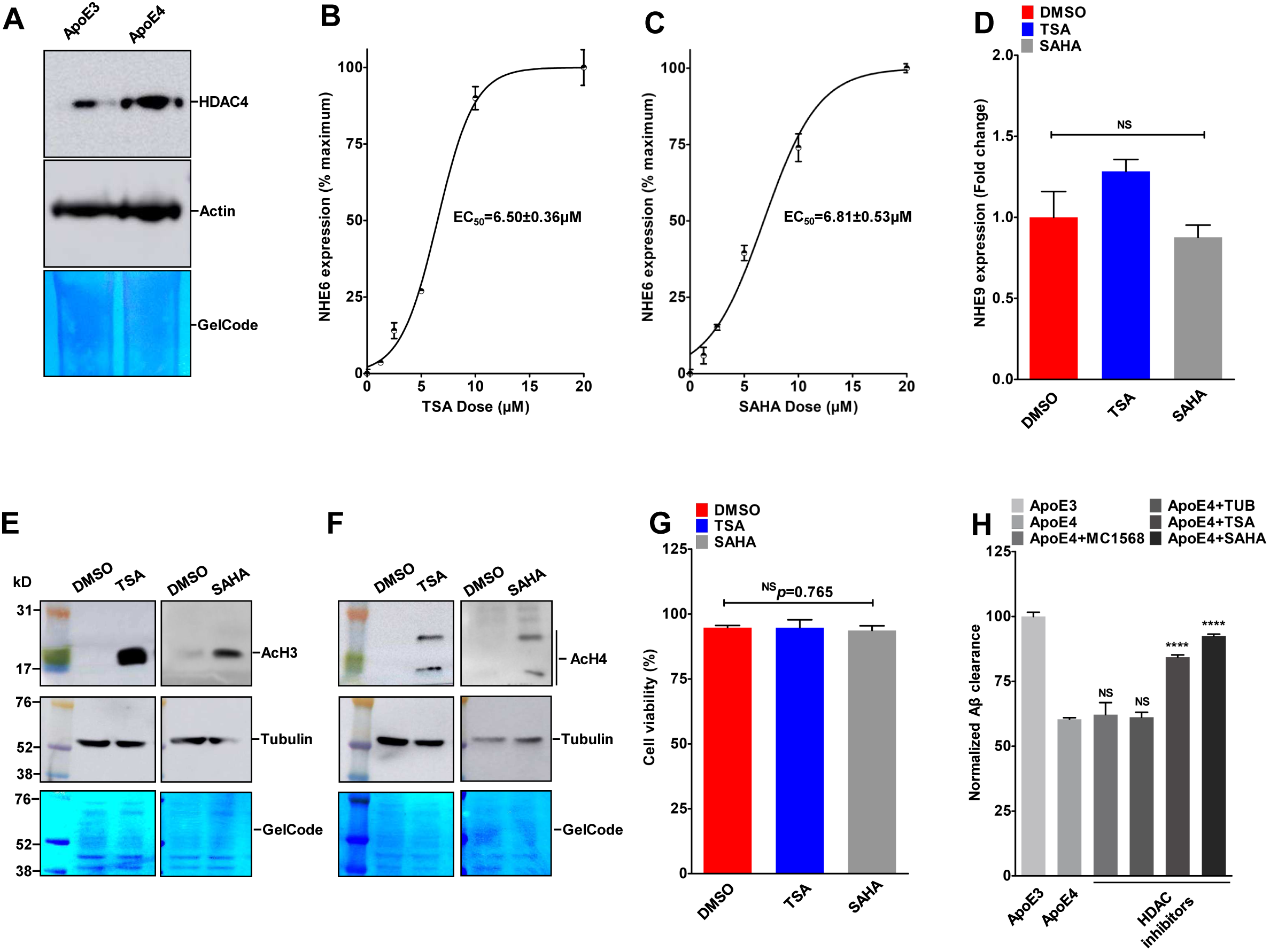
HDAC inhibitors restore NHE6 levels and correct Aβ clearance deficits in ApoE4 astrocytes. Related to Fig 5. A. Western blot of nuclear fractions for HDAC4 levels in ApoE4 astrocytes relative to ApoE3. As loading control, transferred proteins were visualized with Gelcode Blue staining and by probing against actin. Higher HDAC4 in the nuclear fraction is suggestive of increased nuclear translocation of HDACs in ApoE4 cells. (B-C) Dose response relationship to determine the potential of increasing dosage (1.25–20μM) of TSA (B) and SAHA (C) to enhance expression of NHE6 in ApoE4 astrocytes. Note characteristic sigmoidal dose-response curves with half-maximal response (EC50) of 6.50±0.36μM and 6.81±0.53μM for TSA and SAHA, respectively. D. qPCR analysis revealed no significant changes in NHE9 expression with TSA or SAHA treatment of ApoE4 astrocytes. (E-F) Western blot showing acetylation of histone H3 (AcH3) (E) and histone H4 (AcH4) (F) following 60-minutes treatment with TSA and SAHA. As loading control, transferred proteins were visualized with Gelcode Blue staining and by probing against tubulin. G. TSA or SAHA treatment in ApoE4 astrocytes did not affect cell viability measured using trypan blue exclusion. H. Quantitation of measurements of Aβ clearance from FACS analysis of ApoE3 and ApoE4 cells with treated with DMSO and ApoE4 cells with treatment with a panel of HADC inhibitor. Note that broad-spectrum HDAC inhibitors (e.g. TSA and SAHA) that significantly restored NHE6 expression elicited proportionally complete correction of defective Aβ clearance in ApoE4 astrocytes to levels similar (up to 92.4%) to ApoE3 cells. HDAC inhibitors with lower induction of NHE6 expression (e.g. MC1568 and tubacin) conferred minimal changes in Aβ clearance.

## Materials and Methods

### Animals

All procedures were carried out in accordance with The Institutional Animal Care and Use Committee of the University of California, San Francisco and the Johns Hopkins University School of Medicine, Baltimore. The Slc9a6 knockout mice (#005843, strain name B6.129P2-Slc9a6<tm1Dgen) were obtained from Jackson Laboratories. The model was engineered by inserting the LacZ reporter gene, which encodes β-galactosidase into the Slc9a6 genomic locus (Deltagen). In all experiments, male Slc9a6^−/Y^ mice were used as mutants and wild-type male Slc9a6^+/Y^ mice as controls. On average, five mice of each genotype were used in each experiment.

### Aβ clearance assays

Human ApoE isoform-expressing (ApoE3 and ApoE4) astrocyte cells were plated in six-well plates and were grown to confluence. To measure Aβ uptake, cells were washed with serum-free medium (SFM) followed by incubation with 100 nM fluorescently-labeled HiLyte Fluor 647-Aβ40 (#AS-64161, AnaSpec) for various timepoints. Cells were washed with PBS and fixed for confocal imaging using the LSM 700 Confocal microscope (Zeiss), or trypsinized for flow cytometry analysis of ~10,000 cells in biological triplicates using the FACSAria instrument (BD Biosciences). Unstained cells without any exposure to fluorescently-labeled Aβ were used as a control for background fluorescence.

### Aβ assay on mouse brain

The human/rat/mouse β amyloid ELISA kit was from Wako (#294–64701) was used for the estimation of Aβ40 levels in brain homogenates, as per manufacturer’s instructions. Briefly, brains of mice were dissected on ice, weighed and homogenized in ice-old RIPA buffer (PBS+ 1% Triton+ 0.1% SDS+ 0.5% deoxycholate) containing protease inhibitor (Roche). Lysate was centrifuged for 8–10 minutes at 8,000–9,000 rpm and supernatant was collected and used for ELISA. BCA method was used to measure the total protein concentrations. Aβ40 was normalized to total protein concentration in the lysate.

### Endosomal, lysosomal and cytoplasmic pH measurement

Detailed protocols are provided in the Extended Experimental Procedures

### Statistical Analysis

All data were analyzed statistically by Student’s t test, ANOVA and linear regression test using GraphPad Prism. All data are presented as mean ± SD.

## SUPPLEMENTAL INFORMATION

Supplemental Information includes Extended Experimental Procedures and four figures.

### Extended Experimental Procedures

#### Antibodies and Reagents

Mouse monoclonal antibodies used were External epitope Anti-LRP1 (#ab20753, Abcam), Anti-HDAC4 (4A3) (#5392, Cell Signaling Technology), Anti-α-Tubulin (#T9026, Sigma), Anti-β-Actin (#A5441, Sigma), and Anti-GFP (4B10) (#2955, Cell Signaling Technology). Rabbit monoclonal antibodies used were Internal epitope Anti-LRP1 (#ab92544, Abcam), Anti-Na^+^/K^+^-ATPase (#3010, Cell Signaling Technology), Anti-Acetyl-Histone H3 (Lys14) (D4B9) (#7627, Cell Signaling Technology), and Anti-Acetyl-Histone H4 (Lys8) (#2594, Cell Signaling Technology). Monensin (#M5273), KG501 (#70485), Sodium valproate (#P4543), and Sodium butyrate (#B5887) were obtained from Sigma. Clioquinol (#130–26–7) was purchased from Calbiochem. HDAC inhibitors CI994(#A4102), MC1568(#A4094), Tubacin (#A4501), LBH589 (#A8178), TSA (#A8183), and SAHA (#A4084) were from ApexBio Technology.

#### Cell Culture

Human ApoE isoform-expressing (ApoE3 and ApoE4) and ApoE^KO^ immortalized astrocytes (gift from Dr. David M Holtzman, Washington University, St. Louis) were maintained in DME-F12 (Invitrogen) supplemented with 10% fetal bovine serum (FBS) (Invitrogen) and 200μg/ml Geneticin/G418 (Corning Cellgro). Culture conditions were in a 5% CO_2_ incubator at 37°C. Cell viability was measured using the trypan blue exclusion method.

#### Plasmids and transfection

NHE6-GFP and NHE9-GFP were cloned into FUGW-lentiviral vector into the BamHI site. Astrocytes were transfected using lentiviral packaging and expression.

#### Bioinformatics

Mammalian gene expression datasets included in the study were GSE5281, GSE1297, GSE4757, GSE5281, GSE16759, E-MEXP-2280, and GSE15222. Gene Ontology (GO) enrichment analysis was performed using tools provided at the GO web site (http://www.geneontology.org). Pooled analysis was performed using the RevMan program (Nordic Cochrane Centre), as previously described^43^.

#### Transferrin and dextran uptake

Steady state transferrin uptake was measured using flow cytometry and confocal microscopy, as we previously described^28, 43^. Briefly, cells were rinsed and incubated in serum-free medium for 30min, to remove residual transferrin and then incubated with Alexa Fluor 633-Transferrin (#T23362, Thermo Fisher Scientific) (100 μg/ml) at 37°C for 60min. Transferrin uptake was stopped by placing the cells on ice. Excess transferrin was removed by washing with ice-cold serum-free DMEM and PBS, whereas bound transferrin was removed by 2x washing with ice-cold pH 5.0 PBS and pH 7.0 PBS. Cells were fixed with a solution of 4% paraformaldehyde, for confocal imaging using the LSM 700 Confocal microscope (Zeiss), or trypsinized for flow cytometry analysis of ~10,000 cells in biological triplicates using the FACSAria instrument (BD Biosciences). Similarly, dextran uptake was quantified using flow cytometry following 12 hours of incubation with Alexa Fluor 647-Dextran (#D22914, Thermo Fisher Scientific) (10 μg/ml) at 37°C. Unstained cells without any exposure to fluorescently-labeled cargo were used as a control for background fluorescence.

#### Quantitative Real-time PCR

mRNA was extracted from cell cultures using the RNeasy Mini kit (#74104, Qiagen) with DNase I (#10104159001, Roche) treatment, following the manufacturer’s instructions. Complementary DNA was synthesized using the high-Capacity RNA-to-cDNA Kit (#4387406, Applied Biosystems). Quantitative real-time PCR analysis was performed using the 7500 Real-Time PCR System (Applied Biosystems) using Taqman Fast universal PCR Master Mix (#4352042, Applied Biosystems). Taqman gene expression assay probes used in this study are: NHE6, Mm00555445_m1; NHE9, Mm00626012_m1; NHE1, Mm00444270_m1; ATP6V0A1, Mm00444210_m1; LRP1, Mm00464608_m1, ACTB, Mm02619580_g1; and GAPDH, Mm99999915_g1. The Ct (cycle threshold) values were used for all experiments and were first normalized to endogenous control levels by calculating the ΔCt for each sample. Values were then analyzed relative to control, to generate a ΔΔCt value. Fold change was obtained using the equation, expression fold change=2^−ΔΔCt^. Each experiment was repeated three times independently.

#### Immunofluorescence

Cultured cells on polyornithine-coated coverslips were pre-extracted with PHEM buffer (60mM PIPES, 25mM HEPES, 10mM EGTA and 2mM MgCl_2_, pH 6.8) containing 0.025% saponin for 20min, then washed twice for 20min with PHEM buffer containing 0.025% saponin and 8% sucrose. The cells were fixed with a solution of 4% paraformaldehyde (Electron Microscopy Sciences) and 8% sucrose in PBS for 30min at room temperature, and blocked with a solution of 1% BSA and 0.025% saponin in PBS for 1hour. Cells were stained for primary antibodies and Alexa Fluor-conjugated secondary antibodies, DAPI-stained, and mounted onto slides using Dako fluorescent mounting medium and were imaged using a LSM 700 Confocal microscope (Zeiss). Expression of empty GFP vector and NHE6-GFP was detected using the GFP fluorescence. Fractional colocalization was determined from the Pearson’s correlation coefficient, using the JACoP ImageJ plugin that measures the direct overlap of pixels in the confocal section, and represented it as mean ± S.E.

#### Western blot and surface biotinylation

Cells were lysed using 1% Nonidet P-40 (Sigma) supplemented with protease inhibitor cocktail (Roche). Cells were sonicated and then centrifuged for 15min at 14,000 rpm at 4°C. Protein concentration was determined using the BCA assay. Surface biotinylation was performed using Pierce Cell Surface Protein Isolation Kit (#89881, Thermo Fisher Scientific), as per manufacturer’s instructions. Equal amounts of total protein or cell surface protein were separated by polyacrylamide gel (NuPAGE Novex) under reducing conditions and then electrophoretically transferred onto nitrocellulose membranes (Bio-Rad). Ponceau stain or GelCode blue stain (#24590, Thermo Fisher Scientific) was used to confirm protein transfer. Next, the membranes were treated with the blocking buffer containing 5% milk, followed by overnight incubation with primary antibodies and one hour incubation with HRP-conjugated secondary antibodies (GE Healthcare). SuperSignal West Pico substrate was used for detection. Amersham Imager 600 system was used to capture images and densitometric quantification was done using ImageJ software.

#### Endosomal pH measurement

Endosomal pH was measured using flow cytometry, as we previously described ^28, 43^. Briefly, cells were rinsed and incubated in serum-free medium for 30min, to remove residual transferrin and then incubated with pH-sensitive FITC-Transferrin (#T2871, Thermo Fisher Scientific) (75 μg/ml) together with pH non-sensitive Alexafluor 633-Transferrin (#T23362, Thermo Fisher Scientific) (25 μg/ml) at 37°C for 55min. For experiments with GFP tagged NHE6 (wild type and mutants), we used pH-sensitive pHrodo Red-Transferrin (#P35376, Thermo Fisher Scientific), to avoid spectral overlap. Transferrin uptake was stopped by placing the cells on ice. Excess transferrin was removed by washing with ice-cold serum-free DMEM and PBS, whereas bound transferrin was removed by washing with ice-cold pH 5.0 PBS and pH 7.0 PBS. Cells were trypsinized and pH was determined by flow cytometry analysis of ~10,000 cells in biological triplicates using the FACSAria instrument (BD Biosciences). A four-point calibration curve with different pH values (4.5, 5.5, 6.5 and 7.5) was generated using Intracellular pH Calibration Buffer Kit (#P35379, Thermo Fisher Scientific) in the presence of 10μM K^+^/H^+^ ionophore nigericin and 10μM K^+^ ionophore valinomycin.

#### Lysosomal pH measurement

Lysosomal pH was measured as previously described, with modifications^25^. Briefly, cells were rinsed and incubated with pH-sensitive pHrodo-green-Dextran (#P35368, Thermo Fisher Scientific) (5 μg/ml) together with pH non-sensitive Alexa Fluor 647-Dextran (#D22914, Thermo Fisher Scientific) (10 μg/ml) for 12 hours, washed then chased in dye free media for additional 6 hours. Cells were trypsinized and pH was determined by flow cytometry analysis of ~10,000 cells in biological triplicates using the FACSAria instrument (BD Biosciences). A four-point calibration curve with different pH values (3.5, 4.5, 5.5 and 6.5) was generated in the presence of 10μM K^+^/H^+^ ionophore nigericin and 10μM K^+^ ionophore valinomycin.

#### Cytoplasmic pH measurement

Cytoplasmic pH was measured using flow cytometry as we previously described^29^. Briefly, cells were washed and incubated with 1μM BCECF-AM (Thermo Fisher Scientific) for 30min. Cells were trypsinized and pH was determined using flow cytometry analysis of ~10,000 cells in biological triplicates using the FACSAria instrument (BD Biosciences) by excitation at 488nm and emissions filtered through 530 (±15) nm (pH-sensitive green fluorescence) and 616 (±12) nm (pH non-sensitive red fluorescence) filters. A pH calibration curve was generated by preloading cells with 1μM BCECF-AM for 30min followed by incubation in a ‘high K^+^’ HEPES buffer (25mM HEPES, 145mM KCl, 0.8mM MgCl_2_, 1.8mM CaCl_2_, 5.5mM glucose) with pH adjusted to different values (6.6, 7.0, 7.4 and 7.8), and used to prepare a four-point calibration curve in the presence of 10μM K^+^/H^+^ ionophore nigericin.

#### Baculovirus Transduction

For subcellular localization of endocytosed Aβ, we used CellLight-Bacmam 2.0 transduction technology (Molecular probes). Culture medium from ApoE3 astrocyte culture was aspirated, and fresh medium containing CellLight RFP-Bacmam 2.0 reagent targeting early endosome (#C10587, Rab5) or late endosome (#C10589, Rab7), or lysosome (#C10504, Lamp1) was added to reach a final concentration of 30 particles per cell, as suggested by the manufacturer’s instructions. Four hours after transduction 100 nM fluorescently-labeled HiLyte Fluor 647-Aβ40 (#AS-64161, AnaSpec) was added to the media and cultured for an additional 12 hours. Cells were fixed with a solution of 4% paraformaldehyde, DAPI-stained, and mounted onto slides to determine subcellular localization of Aβ by confocal imaging.

#### LRP1 surface labeling

LRP1 surface labeling was performed by immunofluorescence and flow cytometry using a mouse monoclonal antibody against an extracellular epitope of LRP1 (#ab20753, Abcam). For immunofluorescence, live cells were incubated with primary antibody on ice for 1 hour. Cells were then fixed, incubated with secondary antibody, DAPI stained and mounted onto slides to quantify surface LRP1 levels by confocal microscopy. For flow cytometry, live cells were trypsinized and incubated with primary antibody for 30 minutes on ice. Cells were then washed, incubated with secondary antibody, and subjected to flow cytometry analysis of ~5,000 cells in biological triplicates.

#### Aβ surface-binding assay

Ligand surface-binding assay was performed as we previously described, with modifications^29^. Cells on coverslips were rinsed and incubated in serum-free medium for 30min and then exposed to 100 nM fluorescently-labeled HiLyte Fluor 647-Aβ40 (#AS-64161, AnaSpec) in 1% BSA solution on ice for 10min, followed by subsequent washes with PBS at pH 7.0. Cells were then fixed, DAPI stained and mounted onto slides to quantify Aβ surface-binding by confocal microscopy.

#### HDAC nuclear translocation

For quantifying HDAC4 nuclear translation, we immunostained cells using a mouse monoclonal antibody against HDAC4 (#5392, Cell Signaling Technology) and confocal microscopy images were analyzed to measure colocalization between HDAC4 and nuclear stain DAPI. Next, we performed nuclear fractionation of cultured cells using Nuclei isolation kit (#NUC-101, Sigma), as per manufacturer’s instructions. Protein concentration was determined using the BCA assay. Protein from nuclear lysate was resolved using SDS-PAGE and transferred to nitrocellulose membranes. The membranes were then probed with anti-HDAC4 antibody to detect HDAC4 in the nuclear fraction.

